# The crystal structure of the Ca^2+^-ATPase 1 from *Listeria monocytogenes* reveals a pump primed for dephosphorylation

**DOI:** 10.1101/2020.06.23.166462

**Authors:** Sara Basse Hansen, Mateusz Dyla, Caroline Neumann, Esben Meldgaard Hoegh Quistgaard, Jacob Lauwring Andersen, Magnus Kjaergaard, Poul Nissen

## Abstract

Many bacteria export intracellular calcium using active transporters homologous to the sarco/endoplasmic reticulum Ca^2+^-ATPase (SERCA). Here we present three crystal structures of Ca^2+^-ATPase 1 from Listeria monocytogenes (LMCA1). Structures with BeF_3_^-^ mimicking a phosphoenzyme state reveal a closed state, which is intermediate between the outward-open E2P and the proton-occluded E2-P* conformations known for SERCA. It suggests that LMCA1 in the E2P state is pre-organized for dephosphorylation upon Ca^2+^ release, consistent with the rapid dephosphorylation observed in single-molecule studies. An arginine side-chain occupies the position equivalent to calcium binding site I in SERCA, leaving a single Ca^2+^-binding site in LMCA1, corresponding to SERCA site II. Observing no putative transport pathways dedicated to protons, we infer a direct proton counter transport through the Ca^2+^ exchange pathways. The LMCA1 structures provide insight into the evolutionary divergence and conserved features of this important class of ion transporters.

## Introduction

Ca^2+^ regulation is critical for all cells, and therefore also for bacterial cell biology and survival [1]. Active transporters pump Ca^2+^ across the membrane to maintain low intracellular Ca^2+^ concentrations [2]. Mechanistic details of the calcium transport mechanism have been derived mainly for the sarco-endoplasmic reticulum Ca^2+^-ATPase (SERCA) and assumed to extrapolate to other calcium pumps. However, Ca^2+^-ATPases work in a range of different environments across the domains of life, and transport mechanisms must adapt also to specific conditions. To understand how adaptive mechanisms translate sequence variations among Ca^2+^-ATPases to specific functions, detailed structural information of a more diverse pool of Ca^2+^-ATPases is helpful.

The gram-positive bacterium *Listeria monocytogenes* expresses a Ca^2+^-ATPase (LMCA1), which is homologous to mammalian Ca^2+^-ATPases. Soil is the natural habitat of *Listeria*, but they can develop into food borne pathogens, causing listeriosis through infection of the bloodstream, spinal cord membranes and brain. LMCA1 extrudes Ca^2+^ across the bacterial membrane most likely in exchange for a proton [3]. In contrast to mammalian cells, opportunistic bacteria survive in a range of different external environments. LMCA1 thus allows *Listeria* to survive in phagosomal compartments of infected host cells, where Ca^2+^ concentrations can reach millimolar ranges [4]. Furthermore, LMCA1 is part of a complex regulatory network associated with alkaline pH tolerance in the intracellular compartments [5]. LMCA1 is therefore a determinant of *Listeria* intra-cellular Ca^2+^ homeostasis and an important model for both calcium regulation in bacteria and Ca^2+^-ATPase mechanisms.

Ca^2+^-ATPases comprise the P2A and P2B subtypes of the P-type ATPases, which all share a conserved domain structure and key features of their transport mechanisms, involving formation and breakdown of a phosphoenzyme intermediate. The closest mammalian homologue of LMCA1 is SERCA with 34-39% sequence identity, depending on the isoform, while the secretory pathway Ca^2+^-ATPases (SPCA) and plasma-membrane Ca^2+^-ATPases (PMCA) share 34% and 27-30% sequence identity with LMCA1, respectively [3]. Structurally, Ca^2+^-ATPases are characterized by a TM (transmembrane) domain consisting of ten helical segments (M1-10), and a cytosolic headpiece defined by three cytosolic domains: A (actuator), N (nucleotide) and P (phosphorylation) domains (Figure 1A). These domains move relative to each other during the reaction cycle, resulting in transitions between different functional states [6].

**Figure 1.**
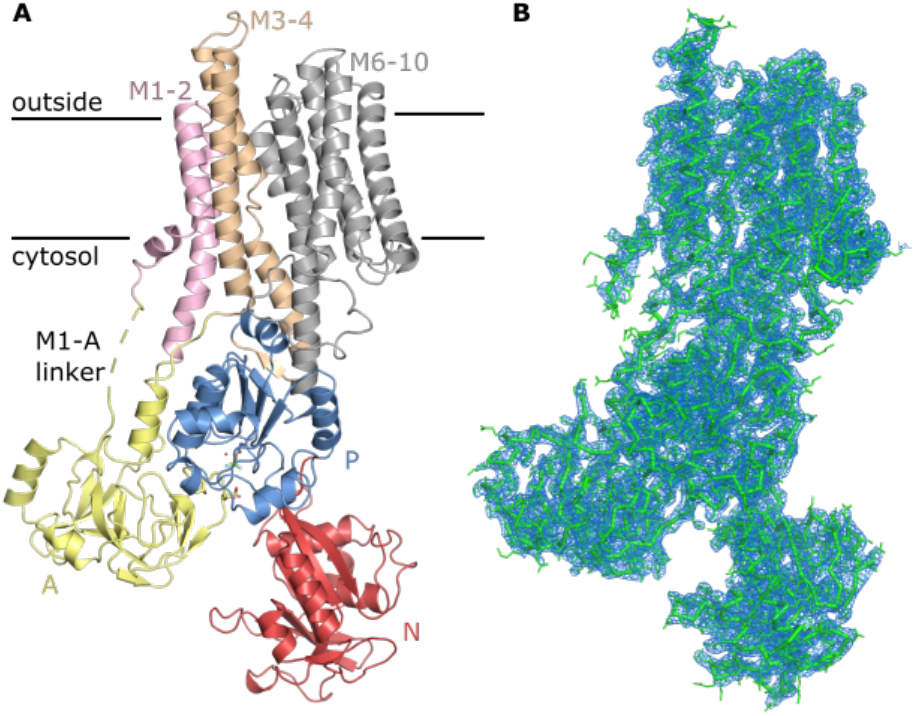
Crystal structure of G_4_E2-BeF_3_^-^ at 3.0 Å. (A) The crystal structure of G_4_ E2-BeF_3_ ^-^ comprises the expected P-type ATPase architecture. Represented as cartoon, it is colored according to the domains. The TGES loop, Asp334 and BeF_3_ ^-^ are shown as sticks. Mg^2+^ and H_2_O are shown as spheres. (B) The 2F_o_-F_c_ map is contoured at 1.5 σ (blue mesh), overlaid with the backbone of G_4_ E2-BeF_3_ ^-^ as ribbon and the side chains as lines (green).

Although SERCA and LMCA1 share functional and mechanistic properties, they are also different. Per molecule of ATP hydrolyzed, SERCA transports 2 Ca^2+^ ions out and counter transports 2-3 protons [7, 8], whereas LMCA1 only transports a single Ca^2+^ ion out and most likely counter transports one proton [3]. Likewise, SPCA and PMCA only transport a single Ca^2+^ ion across the membrane per cycle. Of the two Ca^2+^ binding sites of SERCA, site II is universally conserved among Ca^2+^-ATPases, while site I is replaced with other functionalities in LMCA1 [3], PMCA [9], and SPCA [10]. Homology models and mutational studies of LMCA1 have suggested that an arginine (Arg795) occupies the region corresponding to Ca^2+^ binding site I in SERCA, and may account for a high pH optimum in LMCA1 (pH 8.75-9.5) relative to SERCA (pH ∼7) [3]. Furthermore, kinetic comparisons suggested that the rate-limiting step is phosphorylation for LMCA1, whereas it is the E1P to E2P transition for SERCA [11].

The mechanism of Ca^2+^-ATPases has been described in detail by crystal structures of SERCA trapped by inhibitors at specific intermediate steps of the transport cycle [12]. A schematic diagram summarizing the functional cycle is given in Figure 2A. Overall, ATP hydrolysis drives large structural rearrangements that alter the pump from an inward open (E1) to an outward open state (E2), thereby moving Ca^2+^ across the membrane. In more detail, binding of Ca^2+^ to the E1 state leads to ATP-dependent phosphorylation of a catalytic aspartate in the P domain [13, 14], forming a compact state (E1P). Following phosphoryl transfer, the A domain then undergoes a large rotation, forming the outward-open E2P state [15]. This occurs via a Ca^2+^-occluded E2P state and an ADP releasing step at presumably intermediate positions of the A domain that have not yet been observed in crystal structures, although modelled on the basis of solution X-ray scattering data [16]. Low intracellular ADP concentrations and shielding of the phosphorylated P domain by a conserved TGES loop in the A domain prevent reverse reactions with ADP in the outward-open E2P state, which is crucial for the directionality of the cycle. Negatively charged residues in the transmembrane ion binding sites of Ca^2+^-ATPases are protonated, which allows closure of the outward-open ion pathway. This transition is associated with a small rotation of the A domain around the phosphorylation site that places the glutamate side chain of the TGES loop to activate a water molecule for the dephosphorylation reaction by an in-line nucleophilic attack [17-19]. Dephosphorylation leads to the proton-occluded E2 state stabilized by a hydrophobic cluster that ensures tight association between M2-A and A-M3 linker segments and the P domain with a K^+^ site [20, 21]. Proton release from the ion binding sites to the cytosolic side allows the pump to adopt the inward-open E1 state and again engage in Ca^2+^ binding.

**Figure 2.**
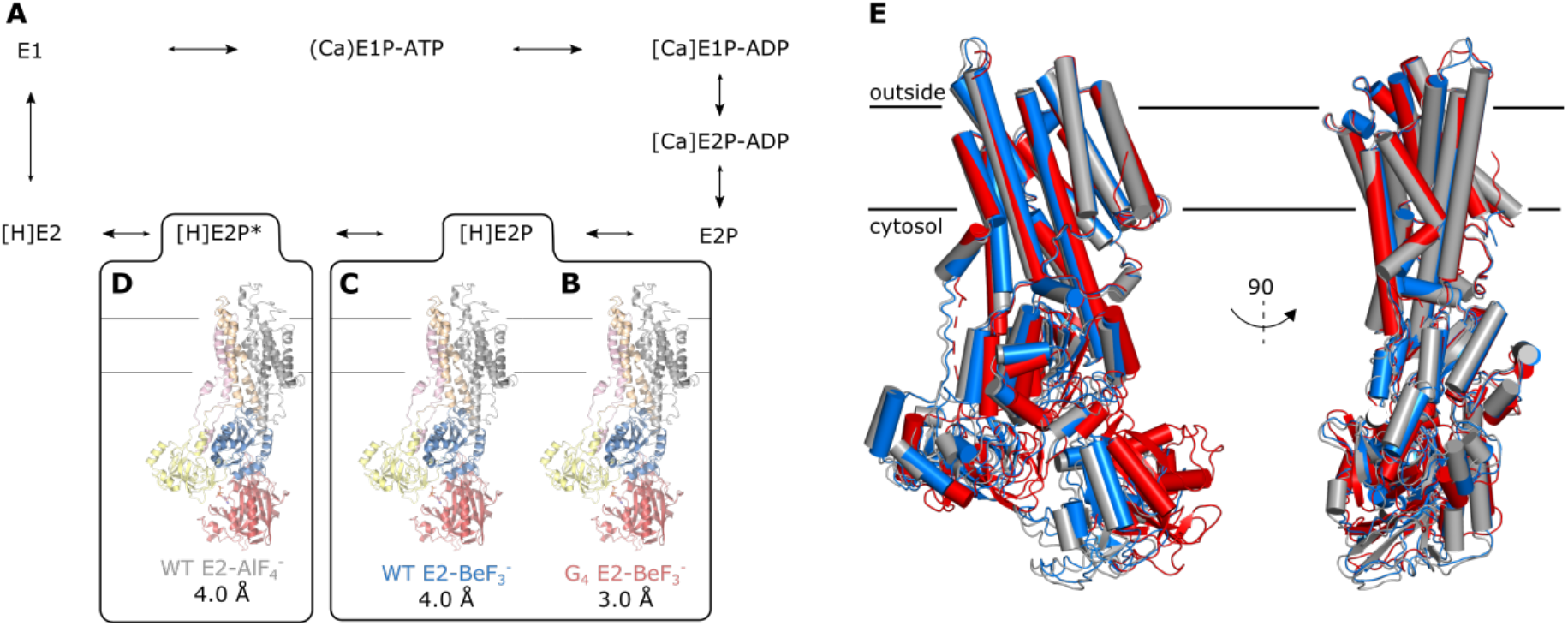
LMCA1 structures adopt proton-occluded E2 states. (A) The reaction cycle of LMCA1 with crystal structures represented by (B) G_4_ E2-BeF_3_ ^-^, (C) WT E2-BeF_3_ ^-^ and (D) WT E2-AlF_4_ ^-^. The structures are colored according to the different domains. (E) An alignment of WT E2-AlF_4_ ^-^ (light grey), WT E2-BeF_3_ ^-^ (blue) and G_4_ E2-BeF_3_ ^-^ (red). All the structures are aligned by transmembrane helices M7-10. A different angle of the cytosolic headpiece relative to the TM domain is observed for the G4 mutant.

Previously, LMCA1 was crystallized in a Ca^2+^-free state similar to the dephosphorylation intermediate (denoted E2-P^*^) using AlF_4_^-^ as a mimic of a pentavalent (trigonal bipyramidal) transition state of dephosphorylation. The structure was partially determined at 4.3 Å resolution [22], but did not permit any detailed analysis. We have also investigated LMCA1 using single-molecule FRET (smFRET) [11, 23]. These data showed that LMCA1 could be paused in E2P states by introduction of a four-glycine insert into the A-M1 linker (G_4_ mutant, identified from SERCA [24]) and most noteworthy, an occluded E2P state preceding Ca^2+^ release was observed. BeF_3_^-^ can be coordinated by the Asp side chain in the phosphorylation site of P-type ATPases, and thereby stabilize a complex mimicking an E2P phosphoenzyme. The smFRET data showed that LMCA1 can be inhibited in mediating dephosphorylation both by forming complex with BeF_3_^-^ and by an E167Q mutant form that targets the TGES motif [11].

We have further explored crystallization of LMCA1 and present here three structures stabilized by metal fluorides: a G4 mutant structure with BeF_3_^-^ at 3.0 Å resolution, and two wild-type structures at 4.0 Å resolution, including a structure with BeF_3_^-^ and an improved structure with AlF_4_^-^.

## Results

To gain new insight into the functional effect of the G_4_ mutant, we crystallized it in the E2-BeF_3_^-^ form and obtained well diffracting crystals at 3.0 Å (Figure 1, Table S1). The asymmetric unit contained 8 molecules related by non-crystallographic symmetry (Figure S1). The refined structure of LMCA1 derived from this crystal form was then used as a molecular replacement model to determine details of LMCA1 dephosphorylation by the structures of wild-type LMCA1 in the E2-BeF_3_^-^ and E2-AlF_4_^-^ forms at 4.0 Å resolution.

### Overall conformation

All three crystal structures show the overall structure expected for a P-type ATPase with three cytosolic domains connected to a membrane domain with 10 transmembrane segments (Figure 1A and 2B-D). Surprisingly, WT E2-BeF_3_^-^ and WT E2-AlF_4_^-^ adopt the same overall conformation, and the configuration of the cytosolic domains is similar to E2-P^*^ like conformations of SERCA in complex with AlF_4_^-^(Figure S2A and Table S2). Like the WT constructs, the E2-BeF_3_^-^ form of the G_4_ mutant adopts a conformation more similar to SERCA E2-AlF_4_^-^ (Table S2), even though the orientation of the cytosolic domains looks more similar to SERCA E2-BeF_3_^-^ (Figure S2B). This can be explained by the position of the cytosolic domains relative to the TM being tilted in G_4_ E2-BeF_3_^-^(Figure 2E). However, the A domain makes different crystal contacts in WT and G_4_ structures (Figure S3), so changes in the domain orientation should be interpreted with caution.

The electron density map reveals no indication of Ca^2+^ bound at the site of the ion binding site in G_4_ E2-BeF_3_^-^ even though the crystallization buffer contained 2 mM Ca^2+^, and all three structures appear to represent occluded states with a closed extracellular pathway and presumably a protonated ion binding site (Figure 2B-D). In the following, we will focus on the structure with the highest resolution (G_4_ E2-BeF_3_^-^) when discussing detailed structural features.

The structures stabilized by BeF_3_^-^ and AlF_4_^-^ inform on transitions associated with extracellular Ca^2+^ release and protonation coupled to dephosphorylation. For the WT E2-BeF_3_^-^ and E2-AlF_4_^-^ structures, the N domain is poorly defined in the electron density maps, and thus likely to be flexible. The asymmetric unit of the G_4_ E2-BeF_3_^-^ crystal contains eight copies of the protein, where four have N domains that are well defined in electron density, while the other four exhibit again weak density for the N domains, indicating flexibility (Figure S4). The flexibility can be related to a minimal role of the N-domain in the dephosphorylation half-cycle. The relative position of the N domain of the G_4_ E2-BeF_3_^-^ structure differs from that of the WT structure, but the P and A domains maintain the same relative configuration, only somewhat tilted relative to the TM domain (Figure 3A). Moreover, the M1-A linker region of the G_4_ mutant appears to be flexible, which uncouples the A domain and the position of the entire cytosolic headpiece relative to the TM domain that is tilted in the G_4_ mutant compared to the WT structures (Figure 2E).

**Figure 3.**
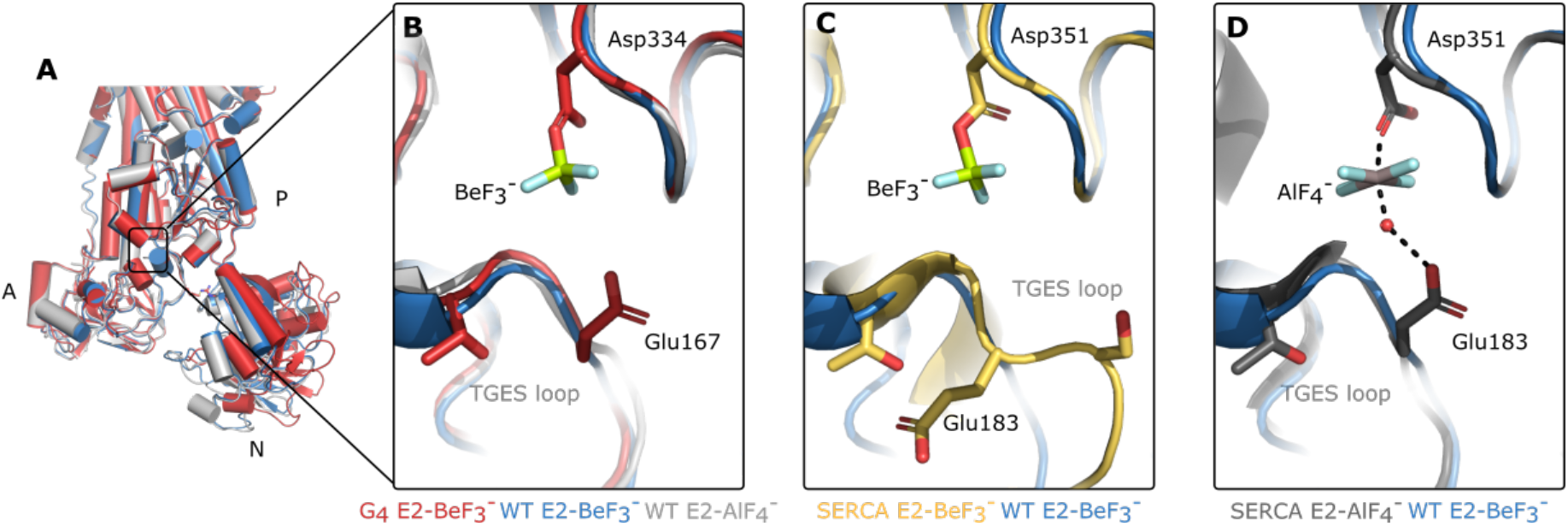
The TGES loop in the E2P state is pre-positioned to catalyze dephosphorylation in LMCA1, but not in SERCA. (A) LMCA1 WT E2-AlF_4_ ^-^ (light grey), WT E2-BeF_3_ ^-^ (blue) and G_4_ E2-BeF_3_ ^-^ (red). (B) Zoom of the TGES loop of (A). The TGES loop of LMCA1 WT E2-BeF_3_ ^-^ aligned with (C) SERCA E2-BeF_3_ ^-^ (pdb: 3b9b) (orange) and (D) SERCA E2-AlF_4_ ^-^ (pdb: 3b9r) (dark grey). All the structures are aligned by the P domain.

#### Interaction between the TGES loop in the A domain and the P domain

The A-P domain interface is critical for dephosphorylation, which is coupled to protonation and occlusion of the extracellular ion exchange pathway. Besides the TGES loop interacting with the phosphorylation site, the interface is also stabilized by Asp186 of the A-domain forming an ionic interaction with Arg598 of the P-domain. Only the G_4_ E2-BeF_3_^-^ structure allows proper refinement of side chain orientations, but similarity of local backbone conformations of the available LMCA1 structures hint at generally preserved interactions in all three structures. In SERCA, the A-P domain interface is connected by several ionic bonds referred to as an ‘electrostatic catch’ centered on Arg198 in the A domain, which is also important for the dephosphorylation rate of SERCA [25]. This residue is not conserved in LMCA1, which indicates that different interactions control LMCA1 dephosphorylation. Despite the fewer interactions defining the A-P domain interface in LMCA1, smFRET experiments have shown that the pump dephosphorylates rapidly, indicating that the required interactions form quickly [11]. With fewer interactions involved, a catalytically competent state may be reached faster in LMCA1.

The two metal fluorides used as phosphate analogues trap LMCA1 in states before and during dephosphorylation. In all three structures presented, the A domain is fully rotated and associated with the P domain in a position that corresponds to the dephosphorylation state of SERCA (E2-P^*^) (Figure 3A). In G_4_ E2-BeF_3_^-^ the TGES loop orients the side chain of Glu167 (Glu183 in SERCA) towards the phosphate analogue, i.e. poised for dephosphorylation (Figure 3B). Position of Glu167 side chain was confirmed by an omit annealed Fo-Fc map (Figure S5A). In SERCA, the calcium-free BeF_3_^-_^ complex adopts an outward-open E2P state, where the TGES loop shields the phosphorylated aspartate and the Glu183 side chain (corresponding to LMCA1 Glu167) points away from the phosphorylation site (Figure 3C). SERCA only adopts an occluded, dephosphorylation state in the AlF_4_^-_^ complex, where Glu183 approaches the phosphorylation site and activates the water molecule that acts as the nucleophile in the hydrolysis (Figure 3D). Since LMCA1 E2-BeF_3_^-^ and E2-AlF_4_^-^ complexes have similar domain arrangements as SERCA E2-AlF_4_^-^, no large-scale conformational changes seem necessary to initiate the phosphatase activity following Ca^2+^ release (Figure 3B). It suggests that LMCA1 adopts a state preactivated for dephosphorylation upon Ca^2+^ release.

### Ion pathways and binding sites

In LMCA1 the Ca^2+^ release pathway is closed in all structures, sealed with an ionic bond between Asp702 (loop M5-6) and Arg261 (M3) and van der Waal interactions. These residues are not conserved in SERCA. The closed extracellular pathway in LMCA1 is further supported by a hydrophobic cluster consisting of Leu107, Met110, Met163, Leu164, Ile215, Val625 and Val646 associating M2, M3 and the P domain at the cytosolic side; this is however conserved in SERCA and represents a characteristic feature of proton-occluded E2 structures [15, 19, 26].

LMCA1 transports a single Ca^2+^ ion and likely counter transports a single proton per hydrolyzed ATP [3]. Alignment of the crystal structures show that the Ca^2+^ binding site of LMCA1 is similar to site II in SERCA (Figure 4). Arg795 on the other hand replaces the Ca^2+^-coordinating Glu908 found in SERCA site I as proposed previously [3]. The other ion-coordinating residue of site I in SERCA (Glu771) [27] is replaced by Ala691 in LMCA1. The short side chain leaves space for the side chain of Arg795 to extend into the core of the LMCA1 TM domain and in effect substitute for Ca^2+^ at the site corresponding to site I in SERCA.

**Figure 4.**
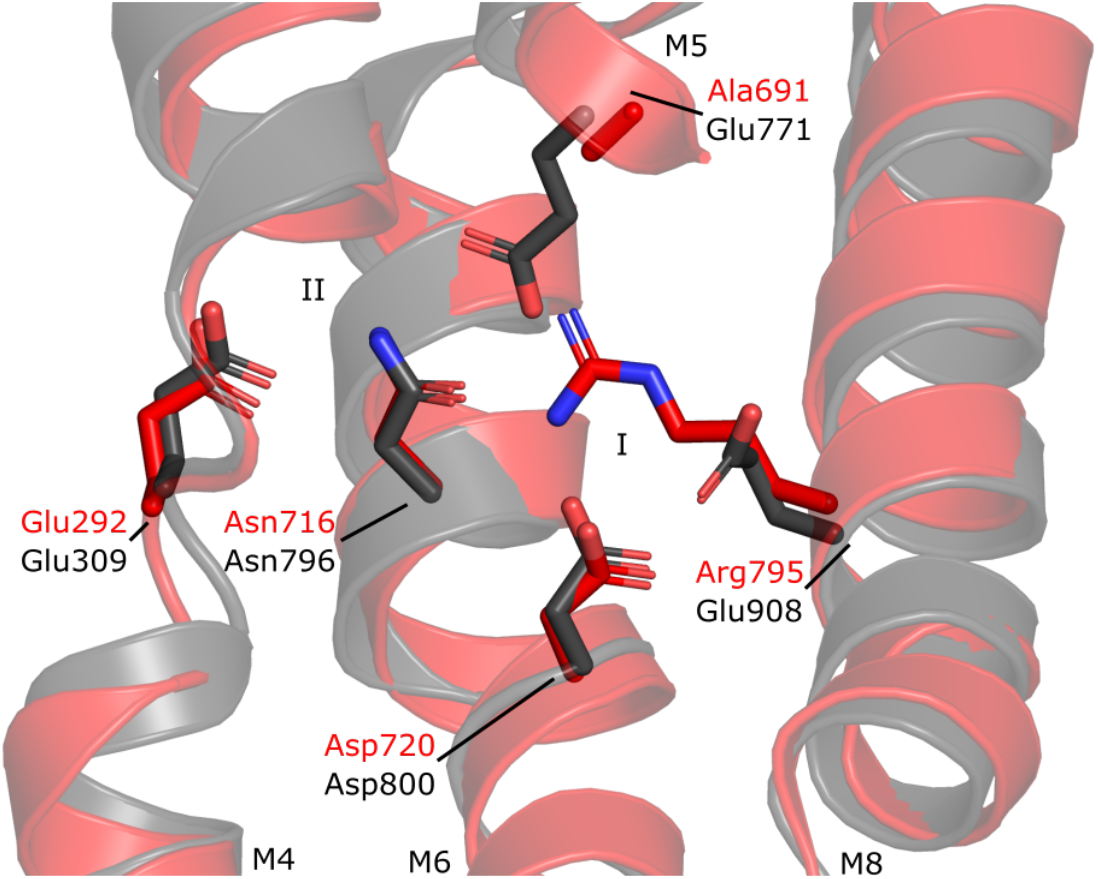
Binding site I of SERCA is not conserved in LMCA1. Alignment between LMCA1 G4 E2-BeF_3_ ^-^ (red) and SERCA E2-AlF_4_ ^-^ (pdb: 3b9r) (dark grey). Relevant Ca^2+^-coordinating residues are shown as sticks. The structures are aligned by the residues shown as sticks. SERCA Ca^2+^ binding site I and II are indicated.

SERCA has two proposed proton pathways: a luminal entry pathway [28] and a C-terminal cytosolic release pathway [29]. These are in addition to the luminal Ca^2+^ release pathway between M1/M2, M3/M4 and M5/M6 [15] and the N-terminal cytosolic Ca^2+^ entry pathway between M1, M2, and M3 [13]. The proposed luminal proton pathway is a narrow water channel lined by hydrophilic residues in M5 and M7, leading protons to binding site I of SERCA [28]. The corresponding part of LMCA1 is mainly hydrophobic, indicating that protons cannot enter through such a pathway.

The C-terminal cytosolic pathway, which is proposed to lead protons from binding site I to the cytosol in SERCA, is also different in LMCA1 (Figure 5). In SERCA, it consists of a narrow hydrated wire between M5, M7, M8 and M10, and the hydrophilic side chains of the residues involved in the water network orient into the cavity (Figure 5A,B) [29]. Although identification of water molecules in LMCA1 is restricted by the resolution of the structures, the nature of the corresponding residues lining the pathway can indicate if a hydrated wire is also present in LMCA1. In fact, the majority of water coordinating residues in the cytosolic pathway in SERCA are different in LMCA1 (Figure 5C,E). For example, Tyr837, Asn914 and Asn981 in SERCA correspond to Gly761, Ala801 and Met868 in LMCA1. The pathway lacks almost entirely polar groups to make a tight interaction network with water molecules. One exception is Arg760 in M7 that extends into the pathway with the aliphatic part occupying the space corresponding to the hydrated cavity in SERCA (Figure 5D). Essentially, M7 is shifted compared to SERCA and Arg760 sterically blocks the pathway in LMCA1. In SERCA, the equivalent position in M7 is Arg836 (Figure 5D). However, this arginine surrounds and coordinates the water molecules in the cavity. Altogether, LMCA1 does not appear to have a continuous water wire that can lead protons from the Ca^2+^ site to the cytosol.

**Figure 5.**
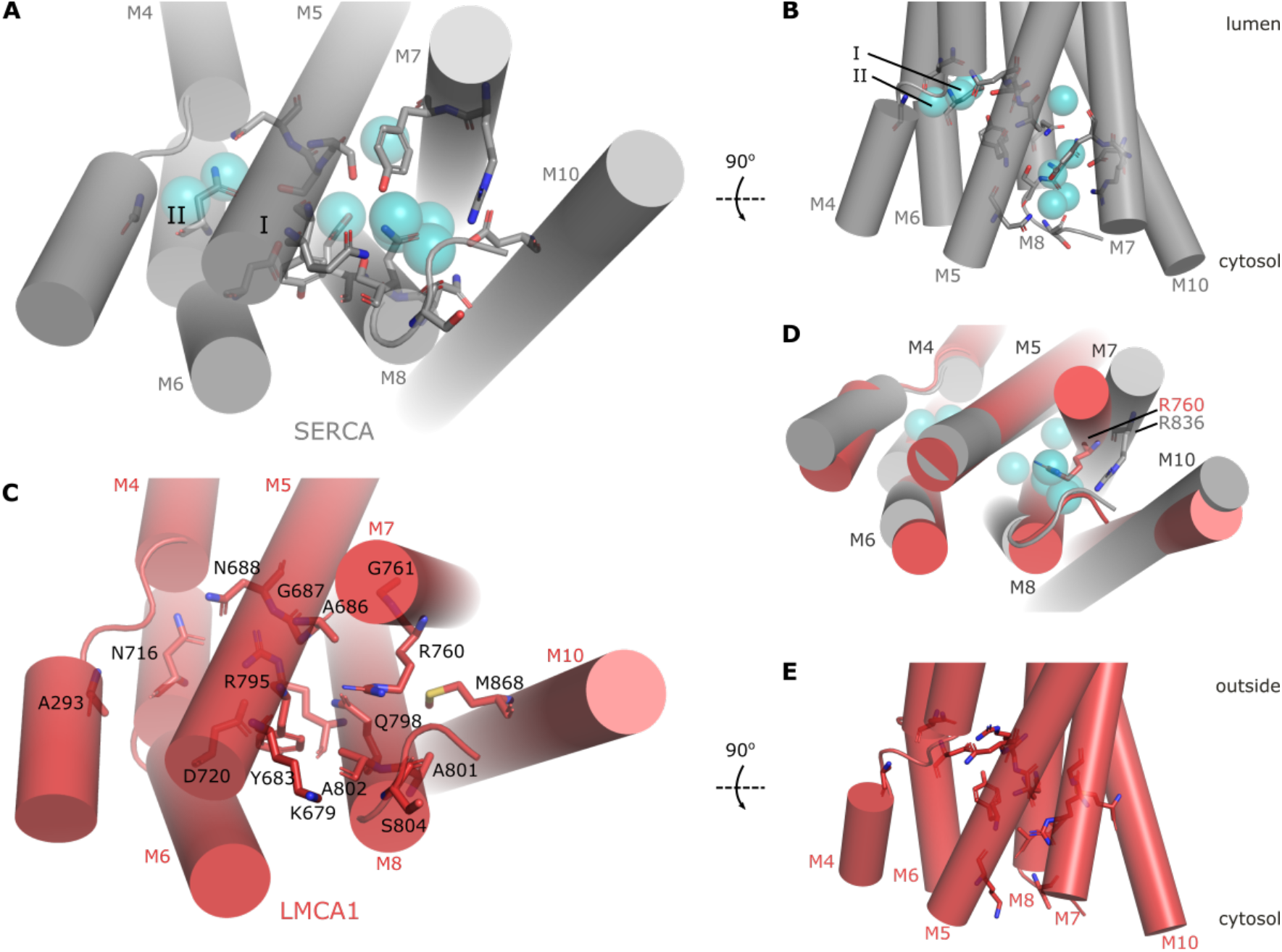
The C-terminal cytosolic pathway leading to site I in SERCA is different in LMCA1. (A, B) G_4_ E2-BeF_3_ ^-^ (red), many of the residues lining the cytosolic cavity are small and hydrophobic. (C, D) In SERCA E2-AlF_4_ ^-^ (pdb: 3n5k) (grey), the C-terminal pathway forms a continuous hydrophilic cavity that leads to ion binding site I. (E) Alignment of LMCA1 and SERCA shows that M7 is shifted and Arg760 extends into the cavity in LMCA1, whereas R836 coordinates the water molecules in SERCA. Water molecules are shown as blue spheres. Only M4-8 and M10 are shown, and the cytosolic domains are omitted for clarity. Residues interacting with water in SERCA E2-AlF ^-^ are shown as sticks, as well as the corresponding residues in G_4_ E2-BeF_3_ ^-^. SERCA Ca^2+^ binding site I and II are indicated.

### Transport site protonation

Ca^2+^ binding sites in Ca^2+^-ATPases are also sites for proton counter-transport by protonation of glutamate and aspartate residues involved in Ca^2+^-coordination. In SERCA, the two glutamates (Glu908 and Glu771) in site I are assumed to be protonated from the lumen in the Ca^2+^ free states and shuttle proton across the membrane in the E2-E1 transition [30]. These proton-transporting side chains are not present in LMCA1, so how are protons then counter-transported? Proton counter transport is observed in LMCA1 [3] and low pH stabilizes the E2 states of LMCA1 [11], suggesting that the ion binding site is indeed protonated in the Ca^2+^-free state. Unless at very high resolution, crystal structures do not reveal protonation directly, but the structure of the ion binding site leaves only two potential candidates that can be reversibly protonated: Glu292 and Asp720 (corresponding to Glu309 and Asp800 in SERCA). To investigate the protonation state of these residues, we estimated p*K*_a_ values based on the G_4_ E2-BeF_3_^-^ crystal structure using PROPKA [31]. The p*K*_a_ values of Glu292 and Asp720 are estimated at 8.9 and 6.2, respectively, which suggests that Asp720 remains negatively charged in the E2P state, whereas Glu292 becomes protonated and therefore likely is the site for proton counter-transport in LMCA1.

## Discussion

We have determined three crystal structures of LMCA1 in E2-BeF_3_^-^ and E2-AlF_4_^-^ bound forms representing the E2P and E2-P^*^ transition state of dephosphorylation. In all cases the TGES loop is pre-positioned for dephosphorylation and the ion binding sites in the TM domain appear to be buried with an occluded proton. It should be noted that occlusion has a slightly different meaning for protons than e.g. Ca^2+^, as it is difficult to experimentally demonstrate proton occlusion. Furthermore, a crystal structure only provides a single snapshot of a protein and it is likely that protein dynamics will allow proton access to a certain extent. In the following, we describe a state as proton occluded, if the ion binding site is closed to direct solvent access.

The structures reveal the architecture of the single Ca^2+^ binding site of LMCA1 in the protonated state and suggest a simpler mechanisms of proton counter-transport compared to SERCA. The structural comparison suggests that residues that constitute the active Ca^2+^ binding site (site II in SERCA) are conserved across a billion years of divergent evolution, but also that LMCA1 and SERCA are individually fine-tuned for different physiological environments. Insertion of four glycines in the linker between the A domain and M1 of LMCA1 causes the enzyme to pause in a Ca^2+^-occluded E2P state [11, 24]. The BeF_3_^-^ stabilized form of this mutant showed no bound Ca^2+^ and represents a proton-occluded pre-state of dephosphorylation, similar to structures of the wild-type enzyme with BeF_3_^-^ and AlF_4_^-^, although with slightly altered angle of the cytosolic domains relative to the TM domain.

### Interdomain interactions account for the fast rate of dephosphorylation

A smFRET study showed that relative rates of partial reactions differ between LMCA1 [11] and SERCA [25]. LMCA1 is rate limited by phosphorylation, dephosphorylates rapidly, and thus predominantly accumulates in E1 states during steady-state pumping [11]. But which structural differences between SERCA and LMCA1 account for the different dephosphorylation rates? Crystal structures of E2-BeF_3_^-^ and E2-AlF_4_^-^ complexes show that LMCA1 and SERCA adopt different conformations when stabilized by the same metal fluorides as phosphate analogues. SERCA E2-BeF_3_^-^ forms an outward-open E2P state with the TGES loop shielding the phosphate analogue. However, the LMCA1 E2-BeF_3_^-^ and E2-AlF_4_^-^ complexes adopt a similar conformation to the proton-occluded SERCA E2-AlF_4_^-^. The LMCA1 structures all have the TGES loop in a position, where the glutamate side chain can orient a water molecule for in-line attack on the phosphorylated aspartate of the P domain (Figures 3B, S5). An outward-open E2P state and a proton-occluded E2-P^*^ intermediate state are likely in rapid equilibrium, but the structural studies point to differences for LMCA1 and SERCA. SERCA changes its conformation to induce dephosphorylation in response to uptake of 2-3 protons through separate proton pathways, and the outward-open E2P state is stabilized by many interactions – these mechanisms can be targeted by regulation. In LMCA1 however, protonation, occlusion and activation of the dephosphorylation site is favored directly upon Ca^2+^ release.

### Binding sites for one or two Ca^2+^

While SERCA transports two Ca^2+^ per cycle, other mammalian P-type ATPases such as a PMCA and SPCA transport only a single Ca^2+^ ion, like LMCA1. A decreased transport stoichiometry increases the driving force for ion transport, but removes regulatory mechanisms enacted by cooperative binding as in SERCA, and likely it represents adaptation to different physiological roles. A low resolution cryo-EM structure of PMCA was determined [9], which allows further comparison between the single Ca^2+^ binding site of mammalian and bacterial Ca^2+^-ATPases. Similar to LMCA1, PMCA has an ion binding site that corresponds to site II in SERCA. In LMCA1, the missing calcium binding site I is filled with a positive charge from an arginine residue at a position corresponding to Glu908 in SERCA - a trait that is shared among a range of bacterial Ca^2+^-ATPases [3]. In PMCA, the corresponding position is occupied by a neutral glutamine [9], while the residue corresponding to Glu771 in SERCA is an alanine for PMCA. No structure has been reported for the SPCA family, but the ion binding site composition can be predicted from homology modeling [32]. No compensating positive charge is found in SPCA, but an aspartic acid occupies the position of the Arg795 in LMCA1, and it may be a unique protonation site. Instead, the calcium site I of SPCA is rendered unfunctional by an alanine in the position of SERCA Glu771, as observed also for LMCA1 and PMCA. This shows that Ca^2+^-ATPases have evolved different mechanisms to alter the transport stoichiometry, and the charge replacement strategy found in LMCA1 is not a unique strategy for stabilizing a non-functional ion binding site.

### Mechanism of proton counter-transport

The altered ion binding site in LMCA1 suggests an altered mechanism of proton counter-transport. Our crystal structures indicate that Glu292 (Glu309 in Ca^2+^ site II of SERCA) has an elevated pK_a_, which is the functional requirement for a protonation site at physiological or even alkaline pH. In all E2-like states of SERCA, a C-terminal water wire connects Glu309 to the cytosol [29], and it is not believed to counter-transport protons. For Glu292 in LMCA1 to participate in counter-transport, the protonation must occur from the extra-cellular side. LMCA1 does not seem to have an additional proton pathway, and protonation must occur directly through the calcium ion exit pathway. LMCA1 thus resembles the P3-type H^+^-ATPases that also seem to feature single entry and exit pathways [33, 34]. This direct mechanism of protonation may relate to the fact that only a single site is involved, thus reducing the need for rapid neutralization of the empty binding site and bifurcated ion pathways [35]. Furthermore, protonation of Glu309 in SERCA is believed to be crucial for closure of the luminal pathway and transition to the proton-occluded state mimicked by E2-AlF_4_^-^ [15]. In an unpredictable, and potentially hostile, external environment, an outward-open state is vulnerable to spurious interactions with compounds. Rapid occlusion and dephosphorylation may also be an evolutionary mechanism of protection for outside-facing pumps.

### The proton-occluded E2P state precedes dephosphorylation

A molecular dynamics study of SERCA proposed that after Ca^2+^ release, the pump transitions into a proton-occluded E2P state with a closed luminal pathway before dephosphorylation [18]. A key prediction of this study was the formation of an intermediate between known crystal structures [18]. This predicted intermediate would correspond to the occluded structure stabilized by BeF_3_^-^ that is observed here. The direct observations of such an intermediate from LMCA1 crystal structures is consistent with these simulations. Such an intermediate may have been observed also earlier in the crystal structure of SERCA in an E2-BeF_3_^-^ form with the inhibitor thapsigargin [36]. This structure features a proton-occluded TM domain and cytosolic domains in the E2P state with the phosphorylation site shielded by the TGES loop. Indeed, the LMCA1 structures determined here are more consistent with the thapsigargin bound SERCA and especially in the orientation of the TM1-2 helices.

### M1-A linker couples the cytosolic domains and TM

The structure of G_4_ E2-BeF_3_^-^ aligns well to WT E2-BeF_3_^-^ in the TM domain, but the cytosolic domains are positioned differently (Figure 2E). In G_4_ E2-BeF_3_^-^, the all the cytosolic domains are tilted compared to WT E2-BeF_3_^-^ and the N domain separates more from the A domain. In fact, we noted two different clusters of conformations in the 8 copies of the asymmetric unit (Figure S1A), where the orientation of the cytosolic domains varies slightly relative to the TM domain (Figure S1B). This suggests that the entire cytosolic headpiece is subject to rigid body movements, and that extension of the A-M1 linker alters the interdomain dynamics.

In SERCA, the G_4_-insertion in the A-M1 linker extends the lifetime of a [Ca_2_]E2-BeF_3_^-^ intermediate state preceding Ca^2+^ release [24], and this state can be entered both through a forward and a reverse reaction. In LMCA1, smFRET data also showed that the G_4_ mutant stabilized the Ca^2+^-occluded E2P state under pumping conditions, albeit only briefly. We crystallized the LMCA1 G_4_ mutant form in the E2-BeF_3_^-^ form in the presence of Ca^2+^, but observed no bound Ca^2+^. The smFRET study of LMCA1 offers a rationale for this, as Ca^2+^ and ADP release was observed to be practically irreversible for LMCA1 [11]. Thus, despite the presence of high Ca^2+^ concentrations, LMCA1 apparently cannot enter this Ca^2+^-occluded E2P intermediate in the reverse direction, and the G_4_ mutant thus crystallized in a Ca^2+^-free form. Unlike the WT complexes, we did not observe density for the central part of the G_4_-extended A-M1 linker in the crystal structure of G_4_ E2-BeF_3_^-^, showing that indeed it gains flexibility and probably weakens mechano-chemical coupling.

## Conclusion

Ca^2+^-ATPases are ubiquitous through-out all domains of life. With structures of a prokaryotic Ca^2+^-ATPase, we illustrate how ATPases have diverged mechanistically and structurally through-out evolution. We observe an intermediate of dephosphorylation that was predicted earlier by molecular dynamics simulations. Future studies should illuminate mechanistic differences of single-site Ca^2+^-ATPases to SERCA-type pumps with cooperative sites.

## Materials and Methods

### Mutagenesis

G_4_-LMCA1: Four glycines between residue K44 and D45 in LMCA1-pET-22b were introduced using the QuickChange mutagenesis kit (Agilent Technologies) and verified by Sanger sequencing (Eurofins, MWG).

### Expression and purification

LMCA1-pET-22b or G_4_-LMCA1-pET-22b, containing a ten-histidine tag and a Tobacco Etch Virus (TEV) protease site in the C-terminus was expressed and purified according to the protocol described in [3]. The solubilized membranes were either applied to a pre-packed HisTrap HP column (GE Healthcare) or incubated with 1 ml Ni^2+^ slurry (Ni-sepharose 6 Fast Flow, GE Healthcare) per 1 L expression medium for 1 hour, which was then packed into an XK-16 column (GE Healthcare). The protein was eluted with 150 mM imidazole in buffer C (50 mM Tris-HCl, 200 mM KCl, 20% v/v glycerol, 1 mM MgCl_2_, 5 mM β-mercaptoethanol (BME), 0.25 mg/ml octaethylene glycol monododecyl ether (C_12_E_8_), pH = 7.6), and then digested with TEV (1 mg per 20 mg protein), while dialyzed against buffer C without imidazole (1:100). The digest was applied to a gravity-flow Ni^2+^ column, and the flow-through collected and concentrated to ∼10 mg/mL using Vivaspin (MWCO = 50 kDa). Size-exclusion chromatography (SEC) was then carried out using a Superdex 200 Increase 10/300 GL column (GE healthcare) equilibrated in 100 mM MOPS, 80 mM KCl, 20% v/v glycerol, 3 mM MgCl_2_, 5 mM BME, 0.25 mg/ml C_12_E_8_, pH = 6.8. Finally, the collected LMCA1 was concentrated to ∼10 mg/mL.

### Relipidation, crystallization and data collection

G_4_ E2-BeF_3_^-^ LMCA1 was relipidated following the HiLiDe approach [17]. A 5 mL glass tube was rinsed with a flow of N_2_ gas to remove the O_2_. 0.3 mg 1,2-dioleoyl-*sn*-glycero-3phosphatidylcholine (DOPC) solubilized in CHCl_3_ was added to the glass tube, and the CHCl_3_ was evaporated with N_2_. Subsequently, 100 *μ*L G_4_ E2-LMCA1 was added, and the tube was sealed with parafilm and stirred at 50 rpm using microstirring bars at 4°C overnight. Insoluble material was removed by centrifugation at 190.000 x g for 10 minutes, and the supernatant was supplemented with 2 mM CaCl_2_ and a premix of 0.1 mM BeSO_4_ and 0.5 mM NaF. 1 *μ*L of the protein solution was mixed with 1 *μ*L of reservoir solution (18% PEG2000, 8% glycerol, 8% MPD, 100 mM MgCl_2_ and 50 mM Tris with pH = 7.2) on a cover slip and equilibrated against 500 *μ*L of the reservoir solution using hanging drop vapor diffusion method. It was sealed with immersion oil (Merck) and equilibrated at 19 °C. Crystals were mounted in loops from mother liquid and flash frozen in liquid N_2_. A complete data set was collected on the P13 beamline at PETRA III radiation source of Deutches Elektronen-Synchrotron (DESY) using a PILATUS 6 M detector.

WT E2-BeF_3_^-^ was relipidated using the same approach as for G_4_ E2-BeF_3_^-^. 0.3 mg DOPC and 0.75 mg C_12_E_8_ solubilized in H_2_O were added to 100 *μ*L protein. After centrifugation, LMCA1 was treated with 2 mM ethylene glycol-bis(β-aminoethyl ether)-*N,N,N*′,*N*′-tetraacetic acid (EGTA) and 1 mM BeSO_4_ premixed with 5 mM NaF. 1 *μ*L of protein solution was mixed with 1 *μ*L of modified reservoir solution (7% PEG6000, 3% t-BuOH, 100 mM LiSO_4_, 5 mM BME, 100 mM KCl, 19 mM C_8_E_4_) on a cover slip and equilibrated against 500 *μ*L of reservoir solution (10% PEG6000, 10% glycerol, 3% t-BuOH, 100 mM LiSO_4_, 5 mM BME) using hanging drop vapor diffusion method. It was sealed with immersion oil (Merck) and equilibrated at 19 °C. Crystals were mounted in loops from mother liquid and flash frozen in liquid N_2_. The expression, purification, relipidation, crystallization and data collection of WT E2-AlF_4_^-^ was described in [22].

### Structure determination

Crystal diffraction data were indexed and integrated with *XDS* [39] and scaled with *Aimless* [40]. The structure of G_4_ E2-BeF_3_^-^ was determined by molecular replacement by *Phaser* [41] using a hybrid search homology model consisting of LMCA1 WT E2-AlF_4_^-^ [22], where the N domain of LMCA1 was based on a homology model of rabbit SERCA-BeF_3_^-^ (PDB: 3b9b) [15]. Homology models were made using an online version of Modeller (www.salilab.org/modeller) [37]. A sequence alignment of LMCA1 and SERCA performed in MUSCLE [38] was used as an input file together with the PDB structures of 3b9b and 3b9r for SERCA stabilized with BeF_3_^-^ and AlF_4_^-^, respectively. The structure determination was challenging, since pseudo-symmetry was present (Figure S1). G_4_ E2-BeF_3_^-^ seemed to display orthorhombic symmetry, since the β angle was close to 90° and the self-rotation function at kappa 180° showed three symmetry axes parallel to the crystallographic axes. However, the processing statistics clearly indicated that orthorhombic symmetry was not correct. The space group is the monoclinic P2_1,_ which only exhibits single two-fold symmetry. The self-rotation function at kappa 180° reveals the presence of rotational pseudosymmetry parallel to the crystallographic symmetry axis. A native Patterson function [42] analysis revealed a peak with a 29% height relative to the origin, which indicates the presence of translational pseudosymmetry. The non-crystallographic symmetry (NCS) operators close to true crystallographic symmetry operators made molecular replacement difficult, but it was eventually achieved, revealing an unusual packing of eight molecules in the asymmetric unit (Figure S1A). NCS averaging was used during refinement and the structure was refined at 3.0 Å maximum resolution. Refinement was made challenging by the fact that large fractions of the N-domain were highly disordered in four out of the eight copies in the asymmetric unit (Table S1). The higher resolution of G_4_ E2-BeF_3_^-^ provided a better start model for structure determination of the WT forms and improving refinement with lower resolution data sets.

WT E2-BeF_3_^-^ and E2-AlF4-crystals both exhibit P2_1_2_1_2 symmetry, The structure of WT E2-BeF_3_^-^ was solved by molecular replacement using G_4_ E2-BeF_3_^-^ as the search model in *Phaser* [41] and refined at 4.0 Å. For the WT E2-AlF_4_^-^ structure, earlier obtained diffraction images [22] were reprocessed in *XDS* [39] and scaled with *Aimless* [40]. Molecular replacement in *Phaser* [41] solved the structure using WT E2-BeF_3_^-^ as a search model. Refinement of all of the structures was done in *PHENIX* [43] and model building and analyses were performed in *Coot* [44].

## Acknowledgements

This work has been supported by grants to P.N. from the Lundbeck Foundation (DANDRITE-R248-2016-2518) and the Independent Research Fund Denmark (FNU) (7014-00328B). S.B.H. was supported by a Ph.D. fellowship from the Boehringer Ingelheims Fond. The authors are grateful for early target identification by J. Preben Morth and the technical assistance of Anna Marie Nielsen and Tanja Klymchuk. We further thank the staff at the EMBL beamline P13 at the PETRA3 synchrotron in Hamburg for access and technical support.

