## Supplemental information for "The crystal structure of the Ca^2+^-ATPase 1 from *Listeria monocytogenes* reveals a pump primed for dephosphorylation"

**Table S1. Data collection and model refinement statistics.** Values in brackets are the highest resolution shell.

| | LMCA1_G$_4$ ($BeF_3^-$) | LMCA1 ($BeF_3^-$) | LMCA1 ($AlF_4^-$) |
|---|---|---|---|
| **Data collection** | | | |
| Beamline | P13 Petra III, DESY | P13 Petra III, DESY | X06SA, SLS |
| Wavelength (Å) | 0.97625 | 0.9762 | 0.9998 |
| Detector | PILATUS 6M | PILATUS 6M | PILATUS 6M |
| Detector distance (mm) | 620.6 | 831.2 | 670.0 |
| Resolution range (Å) | 50 - 3.0 (3.05 - 3.00) | 58 - 3.8 (4.2 – 4.0) | 73 - 3.8 (4.2 – 4.0) |
| Space group | $P2_1$ | $P2_12_12$ | $P2_12_12$ |
| Unit-cell parameters | *a* = 74 *b* = 189 *c* = 350, β=91.9 | *a* = 69 *b* = 144 *c* = 154 | *a* = 181 *b* = 69 *c* = 124 |
| Total reflections | 542,732 | 40,499 | 57,174 |
| Unique reflections | 184,311 | 13,285 | 13,420 |
| Multiplicity | 2.9 (2.8) | 3.0 (3.2) | 4.3 (4.4) |
| Mean *I*/σ(*I*) | 5.8 (1.0) | 4.1 (1.0) | 7.7 (0.9) |
| Completeness (%) | 96.2 (92.7) | 98.4 (47.3) | 97.6 (47.5) |
| Mn(I) half-set correlation CC1/2 (%) | 98.8 (33.2) | 98.9 (73.9) | 99.6 (33.3) |
| $R_{merge}$* (%) | 14.8 (113) | 16.0 (97.0) | 14.4 (275) |
| $R_{pim}$** (%) | 10.1 (79.1) | 10.8 (64.3) | 7.5 (141) |
| Molecules per ASU | 8 | 1 | 1 |
| **Refinement** | | | |
| Resolution | 50 - 3.0 | 58 – 4.0 | 60 – 4.0 |
| R/$R_{free}$ (%) | 25.5/28.5 | 23.2/28.0 | 30.8/32.7 |
| Amino acid residues + ligands | 7,068, 8 $BeF_3^-$, 8 $Mg^{2+}$, 16 water, 15 DOPC, 7 $C_{12}E_8$ | 879, 1 $BeF_3^-$, 1 $Mg^{2+}$, 2 water | 878, 1 $AlF_4^-$, 1 $Mg^{2+}$, 3 water |
| Ramachandran (%) | | | |
| Favored | 95.7 | 94.5 | 95.9 |
| Outliers | 0.2 | 0.3 | 0.1 |
| r.m.s. bond (Å) | 0.007 | 0.003 | 0.006 |
| r.m.s. angle (°) | 0.981 | 0.612 | 1.027 |
| Average B value (Å$^2$) | 128.8 | 169.3 | 159.9 |
| PDB code | 6zhh | 6zhf | 6zhg |

* $R_{\mathrm{merge}} = \Sigma_h \Sigma_i |I_{hi} - \langle I_h \rangle| / \Sigma_h \Sigma_i \langle I_h \rangle$;

** $R_{\mathrm{pim}} = \Sigma_h \Sigma_i (\frac{1}{n_h - 1})^{1/2} |I_{hi} - \langle I_h \rangle| / \Sigma_h \Sigma_i \langle I_h \rangle$

, where $n_h$ is the multiplicity, $I_{hi}$ is the *i*th intensity of reflection *h* and $\langle I_h \rangle$ is the weighted average intensity for all observations *i* of reflection *h*.

**Table S2. Pairwise comparison of LMCA1 and SERCA crystal structures.**

| | ***Full molecule*** | |
|---|---|---|
| *Molecule A* | *Molecule B* | *$C_\alpha$ RMSD (Å)* |
| LMCA1 WT AlF (6zhg) | LMCA1 G4 BeF (6zhh) | 3.356 (866 atoms) |
| LMCA1 WT BeF (6zhf) | LMCA1 G4 BeF (6zhh) | 2.960 (821 atoms) |
| LMCA1 WT AlF (6zhg) | LMCA1 WT BeF (6zhf) | 1.140 (804 atoms) |
| LMCA1 WT AlF (6zhg) | SERCA AlF (3b9r) | 2.945 (820 atoms) |
| LMCA1 WT AlF (6zhg) | SERCA BeF (3b9b) | 4.594 (819 atoms) |
| LMCA1 WT BeF (6zhf) | SERCA AlF (3b9r) | 3.344 (766 atoms) |
| LMCA1 WT BeF (6zhf) | SERCA BeF (3b9b) | 4.552 (763 atoms) |
| LMCA1 G4 BeF (6zhh) | SERCA AlF (3b9r) | 4.351 (842 atoms) |
| LMCA1 G4 BeF (6zhh) | SERCA BeF (3b9b) | 4.790 (822 atoms) |

| | ***Headpiece only**** | |
|---|---|---|
| *Molecule A* | *Molecule B* | *$C_\alpha$ RMSD (Å)* |
| LMCA1 WT AlF (6zhg) | SERCA AlF (3b9r) | 2.851 (443 atoms) |
| LMCA1 WT AlF (6zhg) | SERCA BeF (3b9b) | 3.895 (447 atoms) |
| LMCA1 WT BeF (6zhf) | SERCA AlF (3b9r) | 2.745 (408 atoms) |
| LMCA1 WT BeF (6zhf) | SERCA BeF (3b9b) | 3.749 (422 atoms) |
| LMCA1 G4 BeF (6zhh) | SERCA AlF (3b9r) | 3.423 (463 atoms) |
| LMCA1 G4 BeF (6zhh) | SERCA BeF (3b9b) | 4.408 (462 atoms) |

*The headpiece is defined as follows:
LMCA1: Residues 1-35 + 112-225 + 314-655.
SERCA: Residues 1-37 + 124-241 + 331-735.

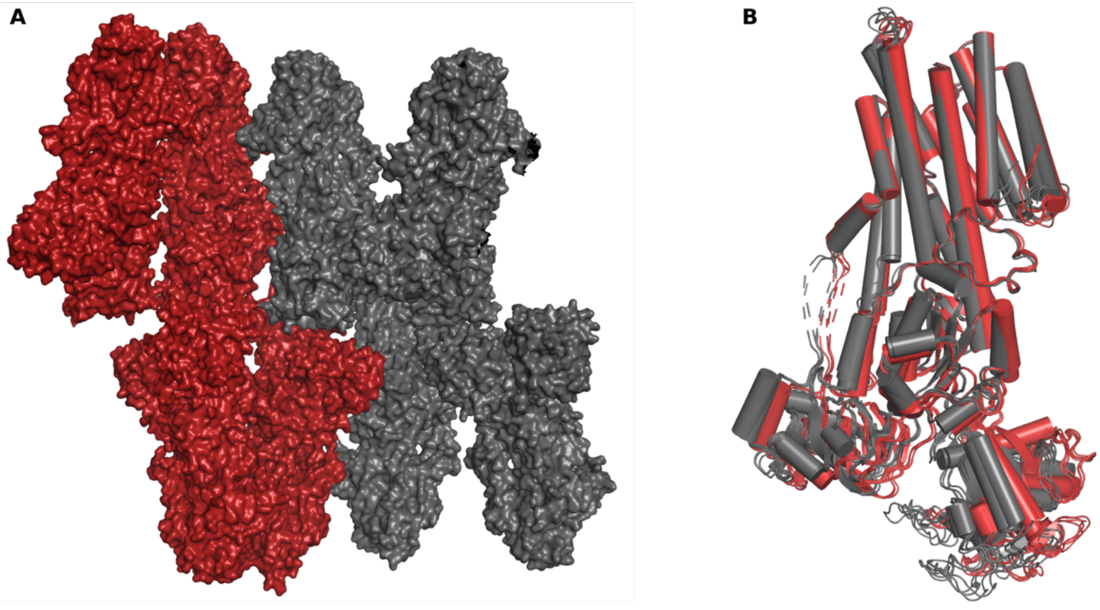

**Figure S1. Packing in the asymmetric unit of the G4 LMCA1 $BeF_3^-$ crystal form.** (A) The asymmetric unit contains two clusters related by non-crystallographic symmetry shown in grey and red. (B) The two clusters show subtle differences in conformations with good alignment of the transmembrane domains and variable orientations of the cytosolic domains. It likely represents an intrinsic flexibility of the fold for the current functional state.

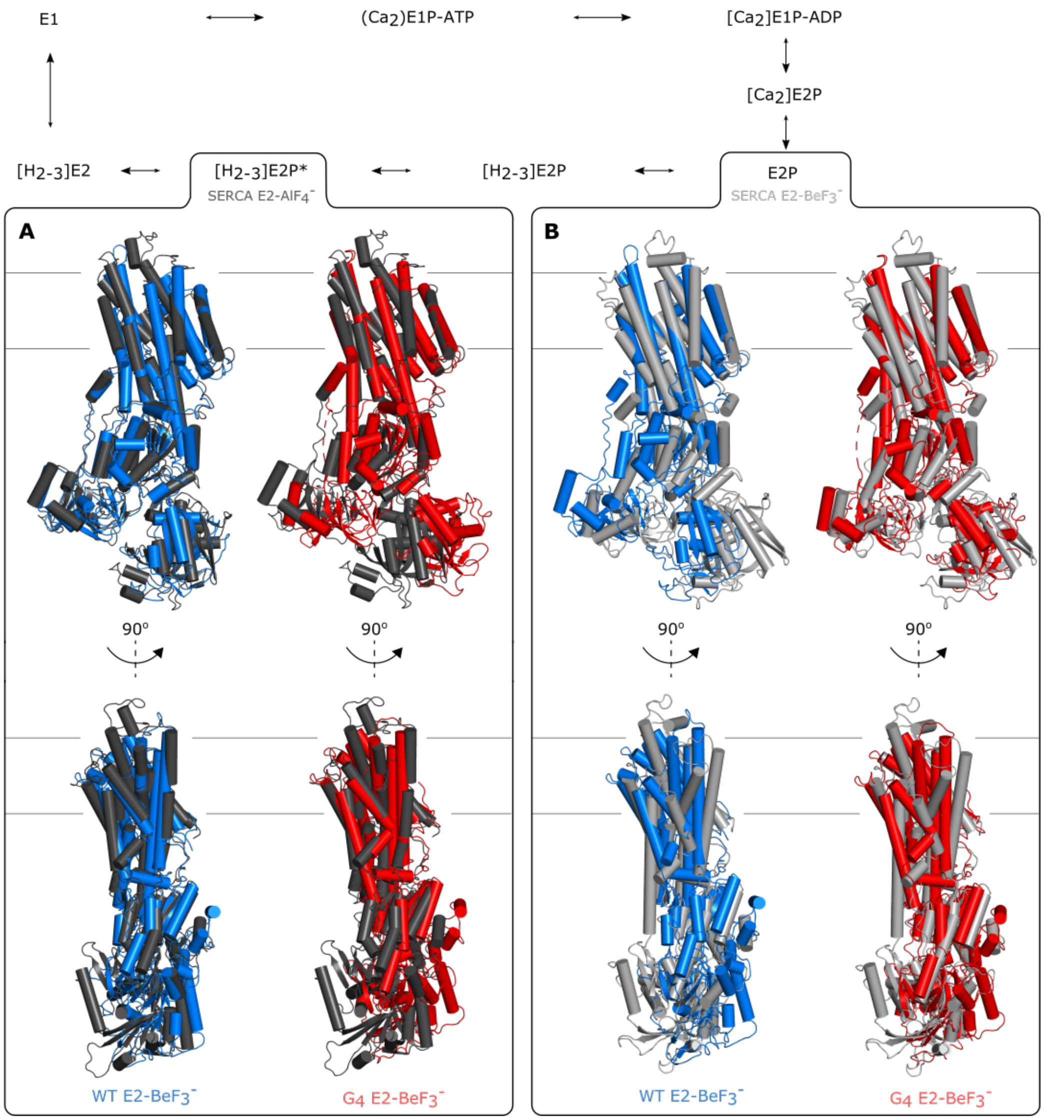

**Figure S2. Comparison of SERCA and LMCA1.** The reaction cycle of SERCA with crystal structures represented as (B) SERCA-$BeF_3^-$ (pdb: 3b9b) (light grey) and (A) SERCA-$AlF_4^-$ (pdb: 3b9r) (dark grey). The LMCA1 WT-$BeF_3^-$ form (blue) has a headpiece orientation more similar to the SERCA E2-$AlF_4^-$ form, whereas the LMCA1 G4-$BeF_3^-$ (red) form have an orientation more similar to SERCA E2-$BeF_3^-$ (RMSD values are given in Table S2). The structures are aligned by transmembrane helices M7-10.

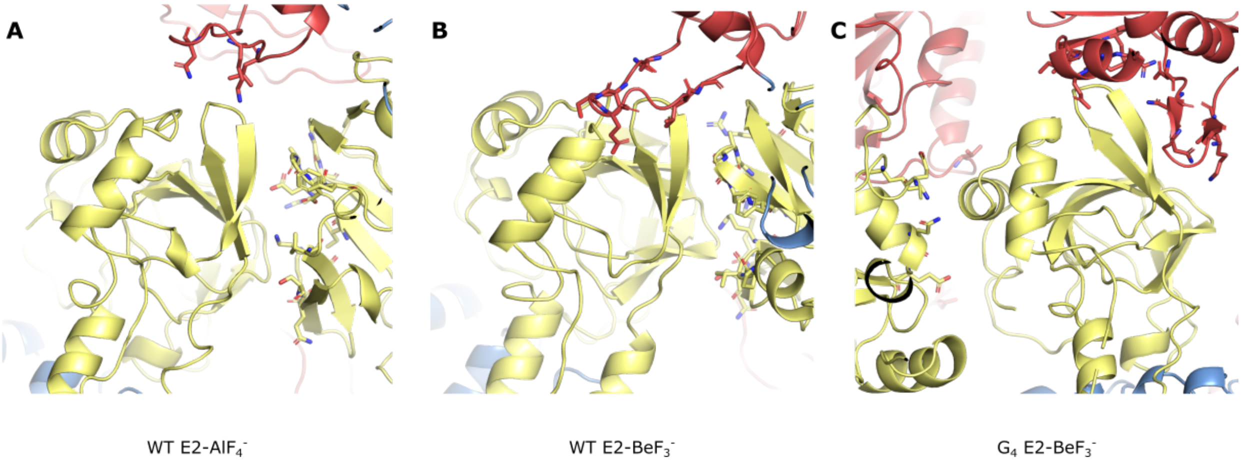

**Figure S3. Crystal packing of the A domain in the three LMCA1 crystal structures.** (A) WT E2-$AlF_4^-$ (B) WT E2-$BeF_3^-$ (C) $G_4$ E2-$BeF_3^-$. Residues in the neighboring molecules within 5 Å of the A domain are shown as sticks. LMCA1 makes different crystal contacts in the G4 structure compared to the WT structures. The models are colored by domain: Yellow: A, red: N, blue: P.

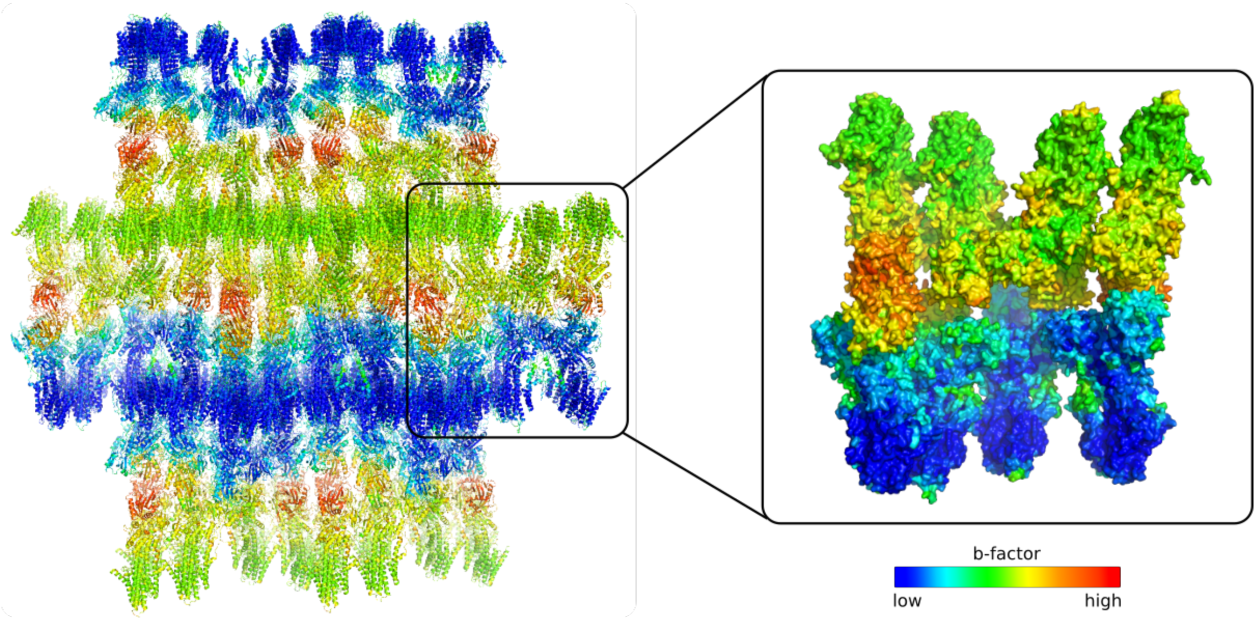

**Figure S4. Variation in structural quality in the asymmetric unit**. B-factors are mapped onto the asymmetric unit ranging from low (blue) to high (red). The B-factors show an uneven distribution, where the packing is better ordered in one layer (blue) compared to the other (green/yellow/orange). The overall packing and crystal contacts in the layers explain the slightly higher R factors than expected for a structure at 3 Å resolution with 61% solvent.

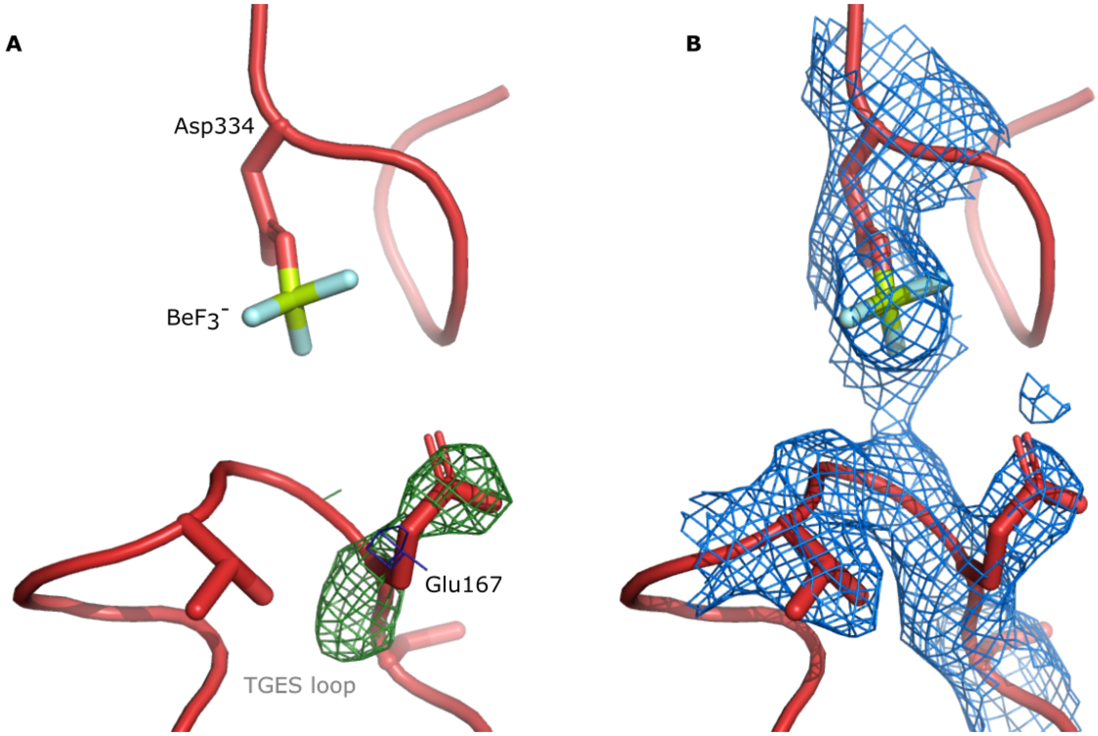

**Figure S5. Orientation of E167 in $G_4$ E2-$BeF_3^-$.** (A) Omit annealed $F_o$-$F_c$ map of Glu167. The map is contoured at -2.5 σ (dark blue mesh) and 2.5 σ (green mesh). (B) $2F_o$-$F_c$ map of Asp334, $BeF_3^-$ and the TGES loop. The map is contoured at 1.5 σ (blue mesh). Molecule B in $G_4$ E2-$BeF_3^-$ was used to show the orientation of Glu167.
